# Natural variation in protein kinase D modifies alcohol sensitivity in *Caenorhabditis elegans*

**DOI:** 10.1101/2024.06.09.598102

**Authors:** Benjamin L. Clites, Brooke Frohock, Emily J. Koury, Oluwatoyin C. Ogunbi, Maya K. Mastronardo, Bhuvan V. Kanna, Erik C. Andersen, Jonathan T. Pierce

## Abstract

Differences in naïve alcohol sensitivity among individuals are a strong predictor of later-life alcohol use disorders (AUD). However, the genetic basis of alcohol sensitivity (beyond ethanol metabolism) and pharmacological approaches to modulate alcohol sensitivity remain poorly understood. We used a high-throughput behavioral assay to measure acute behavioral sensitivity to alcohol, a model of intoxication, in a genetically diverse set of over 150 wild strains of the nematode *Caenorhabditis elegans*. We performed a genome-wide association study and identified five quantitative trait loci (QTL) that underlie natural variation in alcohol sensitivity. We validated that a variant in the conserved ubiquitin-like domain of a *C. elegans* ortholog of protein kinase D, *dkf-2*, likely underlies the chromosome V QTL. Furthermore, lower alcohol sensitivity, *i.e.* resistance to intoxication, was conferred by *dkf-2* loss-of-function mutations. Protein kinase D might represent a conserved, druggable target to modify alcohol sensitivity with application towards AUD.

**Article summary:** We investigated the genetic basis of differences in alcohol sensitivity, a key predictor of alcohol use disorder. We measured alcohol-induced behavioral changes in over 150 genetically diverse nematode strains. Using a genome-wide association study, which links genetic differences to traits, we identified five genomic regions associated with alcohol sensitivity. We then showed that variation in a gene encoding protein kinase D influences resistance to intoxication. These findings identify a conserved molecular pathway that affects alcohol sensitivity and highlight a potential target for intervention. We show how natural genetic variation can reveal mechanisms underlying complex, human-relevant traits.

## Introduction

Alcohol Use Disorder (AUD) is one of the most common psychiatric diseases in the USA, affecting more than one in 10 US adults (Grant *et al*., 2015; 2017). Many more people bear the burden of alcohol misuse through indirect effects, such as drunk driving, lowered productivity, and a stressed medical system, which together cost the USA about a quarter of a trillion dollars annually (Sacks *et al*., 2015). Despite the devastating effects of this disease, relatively few treatment options are available (Franck & Jayaram-Lindstrom, 2013). It is critical that new and effective ways to reduce AUD by treatment and prevention are developed.

Although research into the molecular mechanisms of alcohol has produced many impactful discoveries, a complementary approach has been to identify the specific genetic variants that contribute to population-wide differences in AUD risk (Schuckit, 2018). Identification of the genetic factors that predict AUD could help to target at-risk individuals with early interventions that curtail the development of alcohol-dependence and also provide actionable insights into the biological foundations of AUD. Major progress towards identifying over 100 genetic variants that are associated with problematic patterns of alcohol consumption has been achieved in recent years (*e.g*., Walters *et al*., (2018), Linner *et al*., (2021), Saunders *et al*., (2022)). However, candidate variants and their associated genes need to be functionally validated.

One of the best predictors of future drinking problems is alcohol sensitivity (Schuckit, 1994; 1996; 1998). People with low naïve alcohol sensitivity need to drink more to feel intoxicated and are up to four times as likely to become alcoholics compared to people with normal or high alcohol sensitivity. Differences in alcohol sensitivity exist before AUDs develop and are highly heritable (Verhulst *et al*., 2015). Genome-wide association studies (GWAS) for traits correlated with alcohol use disorder in human populations, including alcohol sensitivity, have advanced rapidly in recent years. Any validated causal variants would make ideal biomarkers for early intervention and AUD risk management. Further, although the complex phenotype of AUD is difficult to model, the relatively simple but highly predictive trait of alcohol sensitivity can be easily modeled in the nematode *C. elegans*.

*C. elegans* has played a key role in identifying molecules with conserved effects in mammalian alcohol responses. Genetic screens identified the BK-potassium channel (SLO-1) as the major target for ethanol in nematodes (Davies *et al*., 2003), and subsequent work used *C. elegans* to identify the role of the NPY (Davies *et al*., 2004) receptor and SWI/SNF chromatin remodelers in alcohol sensitivity (Mathies *et al*., 2015). Mutations in these genes, which were first identified using *C. elegans*, predictably alter alcohol sensitivity in mice and humans (Mathies *et al*., 2015; Du *et al*., 2005; Robinson & Thiele, 2017). Nearly all studies on alcohol response in *C. elegans* have relied on a single genetic background, the laboratory reference strain N2, except for one recent study that analyzed the alcohol responses of a recombinant inbred line panel derived from crosses of six wild strains (van Wijk *et al*., 2023). By leveraging the largely unexplored genetic diversity of hundreds of *C. elegans* wild strains from around the world, additional genes with important orthologous roles in human alcohol sensitivity with relation to AUD can be identified (Crombie *et al*., 2023).

We performed a GWAS with a collection of 151 *C. elegans* wild strains to identify five quantitative trait loci (QTL) that underlie differences in alcohol sensitivity for egg laying behavior. We next showed that the chromosome V QTL is likely explained by variation in *dkf-2*, a highly conserved ortholog of human Protein Kinase D (PKD). Although PKD’s upstream activator, protein kinase C (PKC), has an established role in cross-species alcohol response (Lee & Messing, 2008), our results are likely the first evidence that PKD similarly affects alcohol sensitivity. These findings highlight the power of harnessing natural genetic diversity in a tractable model organism to identify novel genetic effectors of medically important traits such as alcohol sensitivity.

## Materials and Methods

### Animals

Animals were grown at room temperature (20°C) using nematode growth medium agar plates seeded with OP50 *E. coli* bacterial lawns as previously described (Brenner, 1974). Strains were propagated for at least three generations before behavioral testing, to minimize effects of starvation on development and physiology of the animal or its recent lineage. To prevent mutational drift in our test populations, strains were re-thawed from the original stock every one to two months. Animals were synchronized to day-one adulthood using timed-egg lays.

### Assay for alcohol intoxication sensitivity

Nematode growth medium (NGM) agar (12 mL) was dispensed into a 6 cm diameter Petri dish and allowed to set for 48 hours. Next, plates were seeded with ∼0.6 mL of OP50 *E. coli* 48 hours prior to testing. At the start of each assay, 10 day-one adult worms were picked to an untreated control plate, where they were allowed to lay eggs for one hour. Animals were then transferred to an ethanol-treated plate, where they were again allowed to lay eggs for one hour before removal. For each assay, alcohol intoxication sensitivity was calculated by dividing the number of eggs laid on the ethanol plate by the number of eggs laid on the untreated control plate subtracted from one (alcohol sensitivity = 1 - (# eggs laid on ethanol / # eggs laid on control)). To minimize the influence of unforeseen day-dependent variables, strains were tested in blocks of 18-22 variably interleaved across replicates. Genotype averages were calculated using alcohol sensitivity scores from about 10 individual assays. Ethanol plates were prepared by depositing 280 µL of 200 proof ethanol (Sigma) beneath the agar pad 25 minutes before animals are transferred for behavioral testing (Davis *et al*., 2014). After one hour, this exogenous dose results in an internal ethanol concentration of ∼80 mM in day-one adult nematodes (Alaimo *et al*., 2012; Scott *et al*., 2017). To minimize between-assay variation, all ethanol plates were massed to 17.5-18 g before application of ethanol.

### Locomotion

Radial locomotion was quantified as described previously (Topalidou *et al*., 2017; Nordquist *et al*., 2018). In brief, five to 11 day-one adult worms were placed at the center of a 6 cm plate thinly seeded with *E. coli* OP50. After 10 minutes, the radial distance traveled from the center of the plate was measured.

### Heritability calculation

Broad-sense heritability (H^2^) was calculated using H^2^= V_G_/(V_G_+V_E_), where V_G_ is between strain variance and V_E_ is residual variance. H^2^ was estimated using a linear mixed-effect model (*lme4* (R 4.4.0)) where genotype was a random effect (Shaver *et al*., 2023)

### Determining internal ethanol concentrations

Internal ethanol concentrations were measured as described previously (Alaimo *et al*., 2012). In brief, we used a colorimetric assay that uses alcohol metabolic enzymes (ADH & ALDH) to generate quantitatively sensitive shifts in absorbance at 340 nm (Megazyme). Day-one adult animals from each strain were subjected to the same alcohol sensitivity assay as described above. After one hour on ethanol treatment plates, 200 animals of each strain were picked into 120 µL of distilled water. The resulting dilution factor was calculated using estimates of day-one adult worm volumes by Andrews (2019). To prevent evaporation of ethanol from samples, worms were homogenized on ice using a Pyrex grinder for about two minutes per sample. Samples were centrifuged, and supernatant was extracted using a micropipette. Average ethanol concentrations for each strain were determined against a standard colorimetric absorbance curve.

### GWAS

GWAS was performed using egg-laying behavioral sensitivity to alcohol phenotypes of 151 *C. elegans* wild strains and SNV markers from the publicly available CaeNDR *C. elegans* release 20250625 and NemaScan (Widmayer *et al*., 2022). Fine mapping was performed by analyzing genotype-phenotype relationships for all SNVs in the QTL identified by GWAS, rather than only common SNVs (MAF greater than or equal to 5%).

### Strain selection for QTL validation

NIC1 and PS2025 strains were chosen based first on the fact that they did not share many variants in common in the defined QTL, and second that their alcohol sensitivity phenotypes differed by about one standard deviation (SD). NIC1 (mean=0.463) was a little less than one SD below the average phenotype for the ALT group (0.568; SD=0.176) and PS2025 (mean=0.570) was likewise about one SD below the average phenotype for the REF group (mean=0.692; SD=0.093).

### CRISPR-Cas9 genome editing

Ribonucleoprotein (RNP) complexes for co-CRISPR were used to generate mutants (Farboud & Meyer, 2015; Prior *et al*., 2017; Woo *et al*., 2013). Cas9 protein and sgRNA were incubated together to form an RNP complex *in vitro*. The assembled RNP complex and repair template were then directly injected into the gonad of a *C. elegans* hermaphrodite, generating edits in the germline. For ease of screening for desired transformants, a *dpy-10* gRNA was co-injected with a repair oligonucleotide to create a mutation that causes a dominant Rol phenotype (Arribere *et al*., 2014; Prior *et al*., 2017). After confirmation of desired edits by Sanger sequencing, the *dpy-10* marker allele was crossed out by selecting F_2_ animals lacking the Roller phenotype.

Target and repair sequences for *vx40* mutant:

*dkf-2* crRNA: 5’ UGAGGCUCUACAGUUUAUCA 3’

*dkf-2* ssDNA repair oligo: 5’

GGCCATAAGTGTTTTTTAGTTAACGACAGCTTCCTTGACTTTAACGCGAAGCTCTG

GAAGCTTTACTCCGAGATGTCAAATAGTTCCTCCAGCATTCAAAAATAAGGTTTTTTAAGCTTTTTAAACTAT CTTTAAACTTT 3’

*dpy-10* crRNA: 5’

GCUACCAUAGGCACCACGAG 3’

*dpy-10* ssDNA repair oligo: 5’

CACTTGAACTTCAATACGGCAAGATGAGAATGACTGGAAACCGTACCGCAT

GCGGTGCCTATGGTAGCGGAGCTTCACATGGCTTCAGACCAACAGCCTAT 3’

### CRISPR-Cas12a genome editing

Due to the low GC content surrounding candidate variants, it was difficult to find Cas9 PAM sequences (5’-NGG-3’) near our candidate *dkf-2* SNVs. We instead used Cas12a (PAM: 5’-TTTV-3’) to generate single nucleotide substitutions of ALT or REF variants in reciprocal backgrounds. As with Cas9, Cas12a editing was performed using RNPs and co-CRISPR. Alt-R™ A.s. Cas12a (Cpf1) V3 (IDT) was co-incubated with target and *dpy-10* marker gRNAs to form RNP complexes *in vitro*. RNP complexes and ssDNA repair oligonucleotides were co-injected into the germline. F1 individuals showing the Rol phenotype were singled and allowed to lay eggs overnight.

Marker mutations were selected out by isolating F2 individuals with no visible Dpy or Rol phenotypes. Selected F2 individuals were Sanger sequenced to confirm the presence of SNV replacements. If isolated F2 individuals were heterozygous for the targeted substitution, they were allowed to self, and homozygous F3 progeny were isolated and confirmed by Sanger sequencing. Repair oligonucleotides were designed to include blocking mutations, synonymous substitutions which increase the rate of HDR and allele conversion in Cas12a, by reducing unwanted re-cutting of the target sequence in the genome after repair.

Target and repair sequences for allele conversion at *dkf-2* V:16758168:

crRNA: 5’-TCAAGGAAATTGTAAGTTTT-3’

REF to ALT (A->G) repair oligo:

5’GCGCTTCGAGACCTTCGAAATGAGGCTCTACAGTTcATCAAGGAAgTTGTAAGTTTTTATTCCAAAAAATTCGAAAAATTT-3’

ALT to REF (G->A) repair oligo:

5’GCGCTTCGAGACCTTCGAAATGAGGCTCTACAGTTcATCAAGGAAATTGTAAGTTTTTATTCCAAAAAATTCGAAAAATTT-3’

Target and repair sequences for allele conversion at *dkf-2* V:16764489:

crRNA: 5’-AAAATCTTCAAGCACTTGAA-3’

REF to ALT (C->T) repair oligo:

5’CGTTGCTTCTCCTTCTCTTCGTATACTTTCAGATTcTCAAAATCTTCAAaCACTTGAAACGACGCTTGCTGTTGAAATACGAAGC-3’

ALT to REF (T->C) repair oligo:

5’CGTTGCTTCTCCTTCTCTTCGTATACTTTCAGATTcTCAAAATCTTCAAGCACTTGAAACGACGCTTGCTGTTGAAATACGAAGC-3’

Strains (in order of appearance)

*Figure 3b*

N2 – specifically PD1074 version

JPS1627 – *dkf-2(tm4076)* strain CK600 outcrossed 2x

JPS1349 – N2 strain *dkf-2(vx40)* novel CRISPR deletion variant

RB14668 – N2 strain *dkf-2(ok1704)* deletion variant

*Figure 3c*

JPS1849 –– dkf-1(ok2695) strain RB2037 outcrossed 3x JPS1850 – dkf-1(ok2695);dkf-2(tm4076)

JPS1848 – N2 strain immt-2(tm2366) outcrossed 4x

*Figure 3d*

ECA3728 – novel CRISPR strain NIC1(ALT) strain *dkf-2* #1

ECA3731 – novel CRISPR strain NIC1(ALT) strain *dkf-2* #2

ECA3729 – novel CRISPR strain PS2025(REF) strain *dkf-2* #1

ECA3730 – novel CRISPR strain PS2025(REF) strain *dkf-2* #2 *Figure 4*

JPS1859 – novel CRISPR strain N2; dkf-2 (vxIs41[V:16,758,168 A->G]) #1 JPS1860– novel CRISPR strain N2; dkf-2 (vxIs42[V:16,758,168 A->G]) #2 JPS1861– novel CRISPR strain PS2025; dkf-2 (vxIs43[V: 16,758,168 A->G]) #1 JPS1862– novel CRISPR strain PS2025; dkf-2 (vxIs44[V: 16,758,168 A->G]) #2 JPS1863– novel CRISPR strain N2; dkf-2 (vxIs45[V:16,764,489 C->T]) #1 JPS1864– novel CRISPR strain N2; dkf-2 (vxIs46[V:16,764,489 C->T]) #2 JPS1865– novel CRISPR strain PS2025; dkf-2 (vxIs47[V:16,764,489 C->T]) #1 JPS1866– novel CRISPR strain PS2025; dkf-2 (vxIs48[V:16,764,489 C->T]) #2

### PKD inhibitor assays

Alcohol sensitivity assays were conducted as described above. To ensure drug effects, animals were treated with an exogenous concentration of CID 75563 (2.0 μM) that is high relative to its IC50 for human PKDs (0.18-0.28 μM), but still far below the IC50 for non-target protein kinases (*e.g*., PKC = 10.0 μM; PLK1 = 20.3 μM) (Sharlow *et al*., 2008). Exogenous treatment of *C. elegans* to drugs often requires orders of magnitude higher concentrations than effective internal concentrations (Rand and Johnson, 1995; Burns *et al*., 2010). Animals were pre-treated with CID 755673 (DMSO) for one hour prior to alcohol sensitivity assays. During behavioral assays, both the baseline and alcohol egg-laying plates contained CID 75563 (2.0 μM) and 0.01% DMSO. Treated animals were tested alongside same-day controls comprising plates treated with DMSO to account for solvent effects.

### Statistical comparisons

Data were analyzed using SPSS (v29.0.2.0, IBM) or Excel 2022 (365 MSO, Version 2510 Build 16.0.19328.20190, 64-bit) with standard planned two-sided t- or ANOVA tests and the using the Holm-Bonferroni method to correct for *a priori* planned multiple comparisons method and Bonferroni method for *post hoc* multiple comparisons test.

## Results

### A genome-wide association study of alcohol sensitivity identifies five loci

To calculate heritability for *C. elegans* alcohol sensitivity, we assayed a subset of 12 genetically diverse wild strains for acute behavioral sensitivity to alcohol using egg-laying behavior (Fig. 1a,b). Egg laying, which relies on the coordinated action of neurons in a defined circuit, has been exploited to discover and characterize the function of dozens of conserved molecules (*e.g*., Desai *et al*., 1988; Schafer, 2006; Medrano and Collins, 2023). Alcohol-induced depression of egg-laying behavior has served as a model of intoxication in *C. elegans* (Davies *et al*., 2003; Davis *et al*., 2014). We found that alcohol sensitivity differed in the population and was about 42% heritable, mirroring the heritability of alcohol sensitivity observed in human populations (H^2^ ≈ 0.3-0.5; Verhulst *et al*., 2015). To confirm that egg-laying behavior serves as a general readout for alcohol-induced depression in neuromuscular function as in previous studies (Davies *et al*., 2003; Davis *et al*., 2014), we measured alcohol-induced depression in locomotion for six strains and found a strong correlation between the measures (R^2^ = 0.62, p=0.037; Supplementary Fig. 1). Alcohol sensitivity varied widely across strains (*e.g*., genetically diverse set of 11 wild strains compared to the laboratory-adapted N2 strain in Fig. 1b).

**Figure 1:**
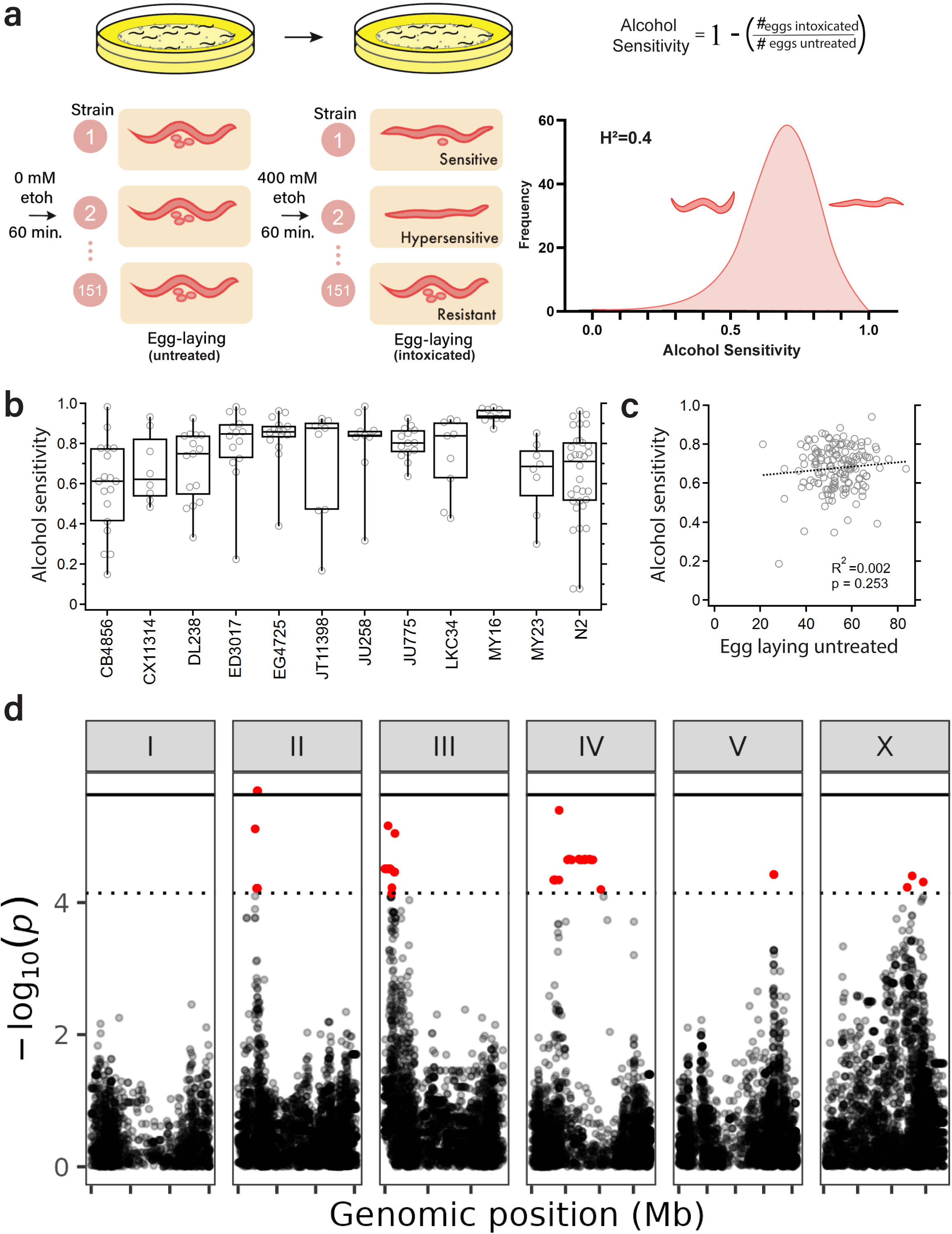
GWAS of natural variation in alcohol sensitivity of 151 *C. elegans* wild strains. **(a)** Overview of high-throughput behavioral screen for naïve alcohol sensitivity in 151 *C. elegans* wild strains. In each assay, basal egg-laying rates were measured using ten clonal adults, and strain average phenotypes were calculated as the average phenotype score across approximately 10 replicates for each genotype. Distribution of alcohol intoxication sensitivity in the population is depicted in top right corner. **(b)** Manhattan plot of GWAS for alcohol sensitivity. Each panel represents a chromosome (I-V & X); each point represents an SNV marker. X-axis represents genomic position (Mb), and y-axis denotes significance (-log_10_(*p*)). Horizontal black lines denote genome-wide significance threshold after correcting for the number of independent association tests: Eigen (dotted) and Bonferroni (solid).

After establishing significant heritable trait variation in the diversity panel, we expanded our assays and measured alcohol sensitivity of 151 wild strains from around the world (Fig. 1a; right; Supplementary Fig. 2). We focused on behavioral sensitivity to alcohol using egg-laying because this trait is easily quantified (number of eggs laid during one hour). The level of alcohol sensitivity could not be simply explained by the untreated level of egg laying because these two measures did not correlate across strains (Fig. 1c). We conducted a GWAS and identified five QTL significantly associated with differences in alcohol sensitivity (Fig. 1d; Supplementary Table 1; Supplementary Fig. 3).

### Candidate genes in the QTL

Our GWAS yielded five QTL that likely contain causal genes that underlie alcohol sensitivity differences. We considered genes with natural variants predicted to alter function and *C. elegans* orthologs of human genes that have been associated with alcohol sensitivity and alcohol use disorder across GWA studies to date. For chromosome II, the most significant variant predicted to affect genes expressed in adult stage *C. elegans* was one in the bidirectional promoter for *K02F6.5* and *K02F6.8* genes that encode potassium channel tetramerization domain (KCTD) proteins. This result is intriguing given that the most significant single gene of large effect conveying low alcohol sensitivity is the BK potassium channel SLO-1 (Davies *et al*., 2003), and two KCTD proteins (6 and 16) have been found by independent human GWAS to be associated with the following alcohol traits: drinks per week, Aspartate aminotransferase platelet ratio index in high alcohol intake, Alcohol use disorder (consumption score), and Fibrosis-4 index in high alcohol intake (Saunders *et al*., 2022; Linnér *et al*., 2019; Kranzler et al;., 2019; Innes *et al*., 2020). Although the BK channel does not appear to rely on KCTD proteins to tetramerize, KCTD affects other potassium channels and has additional roles such as interacting with Gβγ to alter cAMP signaling (Sloan *et al*., 2023). The chromosome II region also included *sup-9*, which is orthologous to human *KCNK3* potassium channel gene that has also been associated with an alcohol trait (Diastolic blood pressure x alcohol consumption (light vs heavy) interaction (2df test)) (Feitosa *et al*., 2018). Because alcohol sensitivity in *C. elegans* correlates with anesthetic sensitivity, the potassium channel *sup-9* is a strong candidate (Morgan *et al*., 2007). Loss of function confers resistance to halothane-induced paralysis, whereas gain of function in its channel component, *sup-10*, causes hypersensitivity to halothane (Singaram *et al*., 2011). For chromosome III, the QTL peak was in *pll-1,* which is an ortholog of human PLCL1 that was associated with alcohol consumption in a meta-analysis of human GWAS (Kapoor *et al*., 2013). The edge of this QTL contains the acetylcholine transporter *snf-6*, which was isolated in the original forward-genetic screen for mutants with low alcohol sensitivity (Davies *et al*., 2003; Kim *et al*., 2004). All of these candidate genes are compelling, but without specific genome-editing experiments to establish causality, they remain candidates.

The QTL on chromosomes IV and X are more difficult to parse. This result might be explained in part by the low recombination rate of the X chromosome leading to QTL with a larger number of linked genes. Because of the hyper-divergent regions (Lee *et al*., 2021) that overlap the QTL on chromosomes II, III, and IV, the gene content is difficult to dissect further. In *C. elegans*, certain regions of the genome contain high levels of genetic variation and include huge amounts of novel genomic content (Lee *et al*., 2021; Moya *et al*., 2025). Although hyper-divergent regions are difficult to dissect, they still could represent true effectors of *C. elegans* alcohol sensitivity.

Given the significance of the five QTL along with the candidates from fine mapping, we chose to pursue the QTL on chromosome V. This QTL encompassed an approximately 1 Mb interval, where the ALT allele predicted significantly lower alcohol sensitivity, *i.e*., resistance to intoxication (Fig. 1b; Fig. 2a). To identify potential causal variants in the chromosome V QTL, all rare and common SNVs in the interval were tested for association with alcohol sensitivity differences, with priority given to alleles with high predicted effects on gene function (*e.g*., nonsynonymous substitutions, early stop codons, splice-site variants, etc.) (Fig. 2b). Fine mapping identified several high-impact variants in the two nearest genes to the peak QTL marker, *dkf-2* (V:16,764,489 C->T) and *immt-2* (V:16,790,476 G->A), that segregated with differences in sensitivity to ethanol intoxication (Supplementary Fig. 4). The gene *dkf-2* encodes a *C. elegans* ortholog of protein kinase D (PKD) (Feng *et al*., 2007). Interestingly, PKDs function downstream of PKC, a known effector of alcohol response (Wallace *et al*., 2007), so we focused our efforts on this candidate gene. Further, deletion of *dkf-2* was shown to restore movement in a TDP-43 model of neurodegeneration (Liachko *et al*., 2014), suggesting an ameliorative effect on neuromuscular depression. These data together suggest that ALT variants in *dkf-2* might underlie the QTL for alcohol sensitivity.

**Figure 2:**
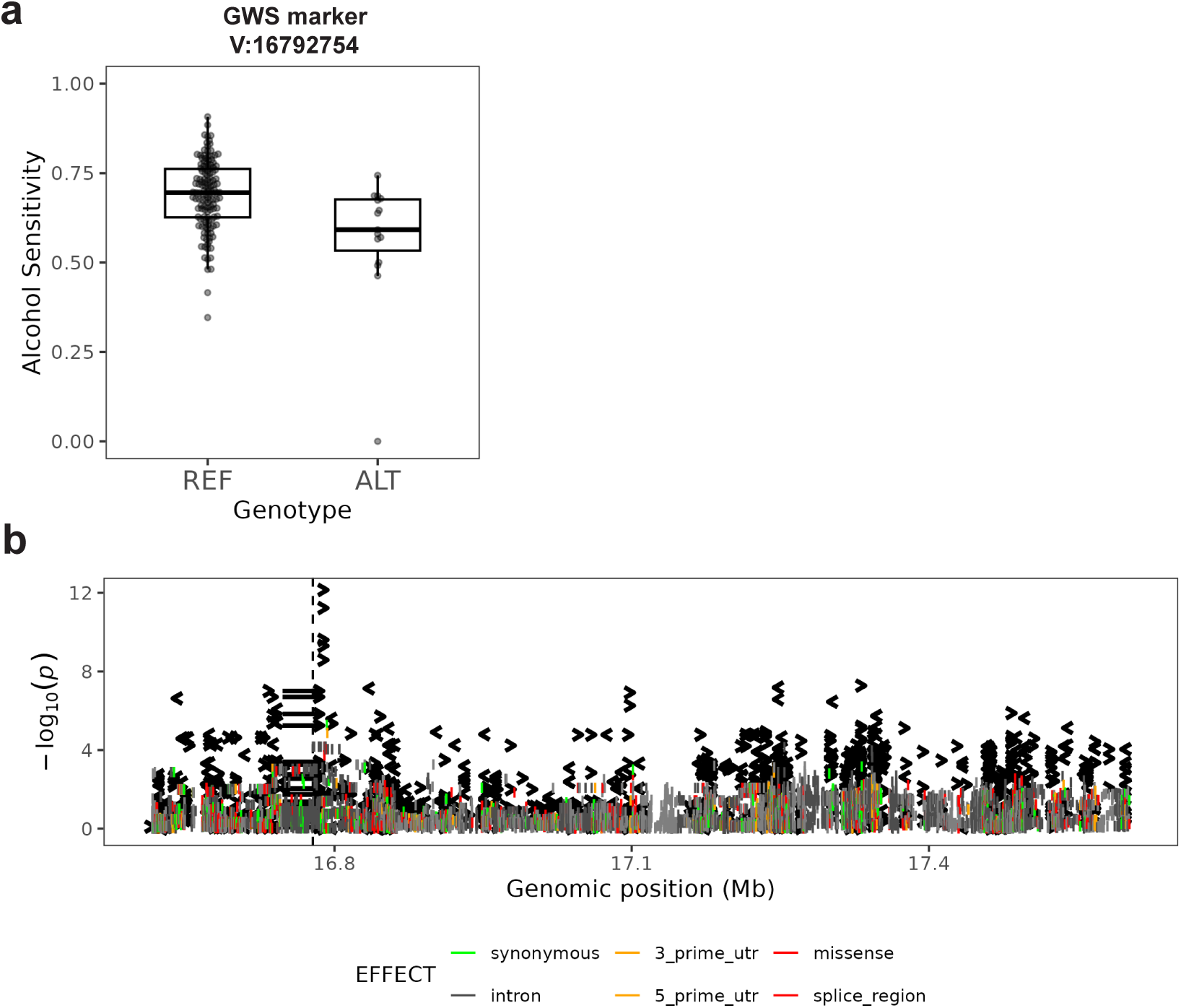
Fine mapping the chromosome V QTL associated with decreased sensitivity. **(a)** Each point in boxplots represents the average alcohol sensitivity of a single strain. Y-axis denotes alcohol intoxication sensitivity, and genotype at SNV is shown along the X-axis (REF or ALT). Average phenotypes of all strains were segregated by their genotype at the denoted allele. **(b)** Results from fine mapping. Each line represents an SNV in the significant interval (V:16,638,047-18,330,166) from GWAS. Y-axis denotes significance. Color fill of each SNV represents predicted variant effect, where red denotes high-impact variants, gray represents low-impact variants, and light gray represents intergenic/linker variants.

### Validating the role of Protein kinase D in alcohol sensitivity

To test whether *dkf-2* represented a causal gene for alcohol sensitivity differences, we assayed *dkf-2* loss-of-function mutants. We obtained two *dkf-2* deletion alleles in the N2 laboratory wild-type background: *ok1704*, a 716 bp deletion in exons 11 and 12, and *tm4076*, a 1,646 bp deletion that interrupts an obligate exon of all six known isoforms (Fig. 3a; Supplementary Fig. 5). We found that strains harboring either of the two deletion alleles had lower alcohol sensitivity relative to the N2 strain (Fig. 3b). We also generated a novel deletion allele, *vx40*, that deletes two exonic bases causing a putative frame-shift loss-of-function allele. This novel deletion also conferred decreased alcohol sensitivity. Interestingly, *vx40* affects only four of the six predicted *dkf-2* isoforms (a, d, e, and f), one fewer than the *ok1704* deletion (a, c, d, e, and f) and two fewer isoforms than the *tm4076* deletion (Hillier *et al*., 2009) (Supplementary Fig. 5). These results provide evidence that loss of *dkf-2* causes low alcohol sensitivity.

**Figure 3:**
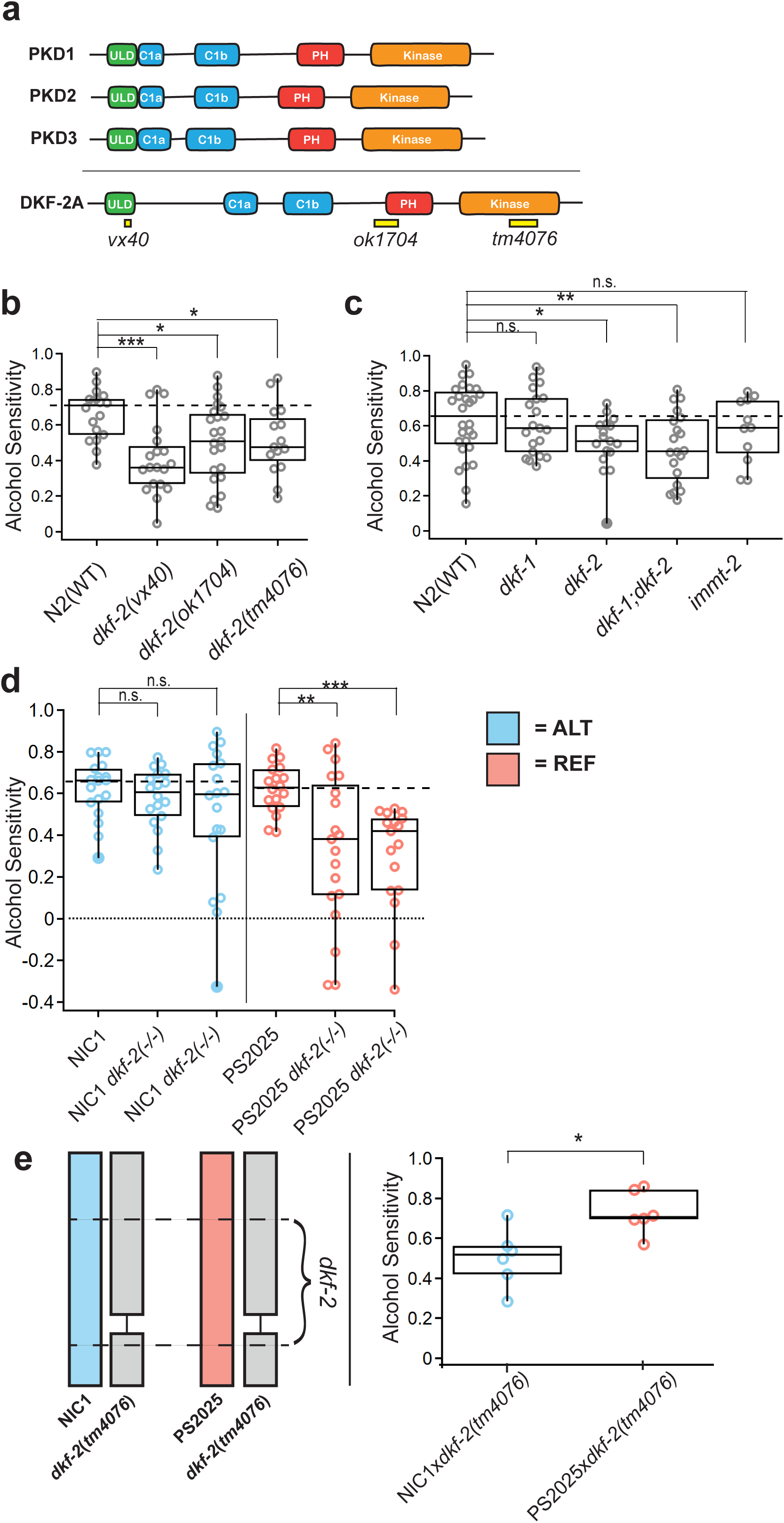
Independent deletion alleles in the highly conserved *C. elegans* ortholog of PKD (*dkf-2*) predictably reduce alcohol sensitivity. **(a)** Protein kinase D is functionally conserved from humans to *C. elegans*. The human gene products (PKD1, PKD2, PKD3) are compared to one isoform of the *C. elegans* ortholog (DKF-2-A). Orthologous functional domains are labeled. **(b)** Alcohol sensitivities of three *dkf-2* deletion alleles (*vx40*, *ok1704*, and *tm4076*) were significantly lower than intoxication sensitivity of *dkf-2* lab wild-type (N2) animals. (n=15; *p*=0.0001; p=0.0104; p=0.0167). **(c)** Alcohol sensitivities of a *dkf-1* and *immt-2* mutants were not significantly different from laboratory-adapted N2 strain, but both *dkf-2* and *dkf-1; dkf-2* double mutants were significantly lower than the N2 strain(n=11-28; p=0.048; p=0.001). **(d)** Deletion mutations in a wild strain (NIC1) harboring candidate hypomorphic allele(s) in *dkf-2* does not cause a reduction in alcohol sensitivity (mean & SEM; n=18; p=0.35; p=0.1573), whereas deletions in a strain without the candidate hypomorphic alleles cause significant reductions in alcohol sensitivity (mean & SEM, n=18; p=0.0001; p=0.0052). **(e)** Strains that are hemizygous for ALT genotype at *dkf-2* are resistant to alcohol, although strains that are hemizygous for the REF genotype at *dkf-2* are sensitive to alcohol (mean & SEM; n=6; p=0.01). NIC1 (ALT) and PS2025 (REF) were mated to a *dkf-2* deletion mutant (*tm4076*). F_1_ progeny from these crosses were assayed for their alcohol sensitivity. Alcohol sensitivity was higher in PS2025 (REF)x*dkf-2*(-/-) F_1_’s relative to NIC1(ALT)x*dkf-2*(-/-) F_1_’s. Planned two-sided t-tests were used for comparisons. Box plots show the median, interquartile range, data spread, and potential outliers for each group.

To seek further context on the role of *dkf-2* in alcohol sensitivity, we also tested *immt-2* and *dkf-1* mutants. *immt-2* is an ortholog of human IMMT (inner membrane mitochondrial protein) which is adjacent to *dkf-2* in the genome (Sup Fig 4). *dkf-1* encodes the only other PKD kinase in *C. elegans*. (Feng et., 2006). We found that the *immt-2* and *dkf-1* mutants displayed indistinguishable sensitivity compared with N2 control (Fig. 3c). In contrast, *dkf-2* single and *dkf-1; dkf-2* double mutants, assayed in parallel, both showed significantly lower sensitivity compared to N2. We conclude that loss of function of *immt-2* or *dkf-1* is unlikely on its own to contribute to reduced alcohol sensitivity.

**Figure 4:**
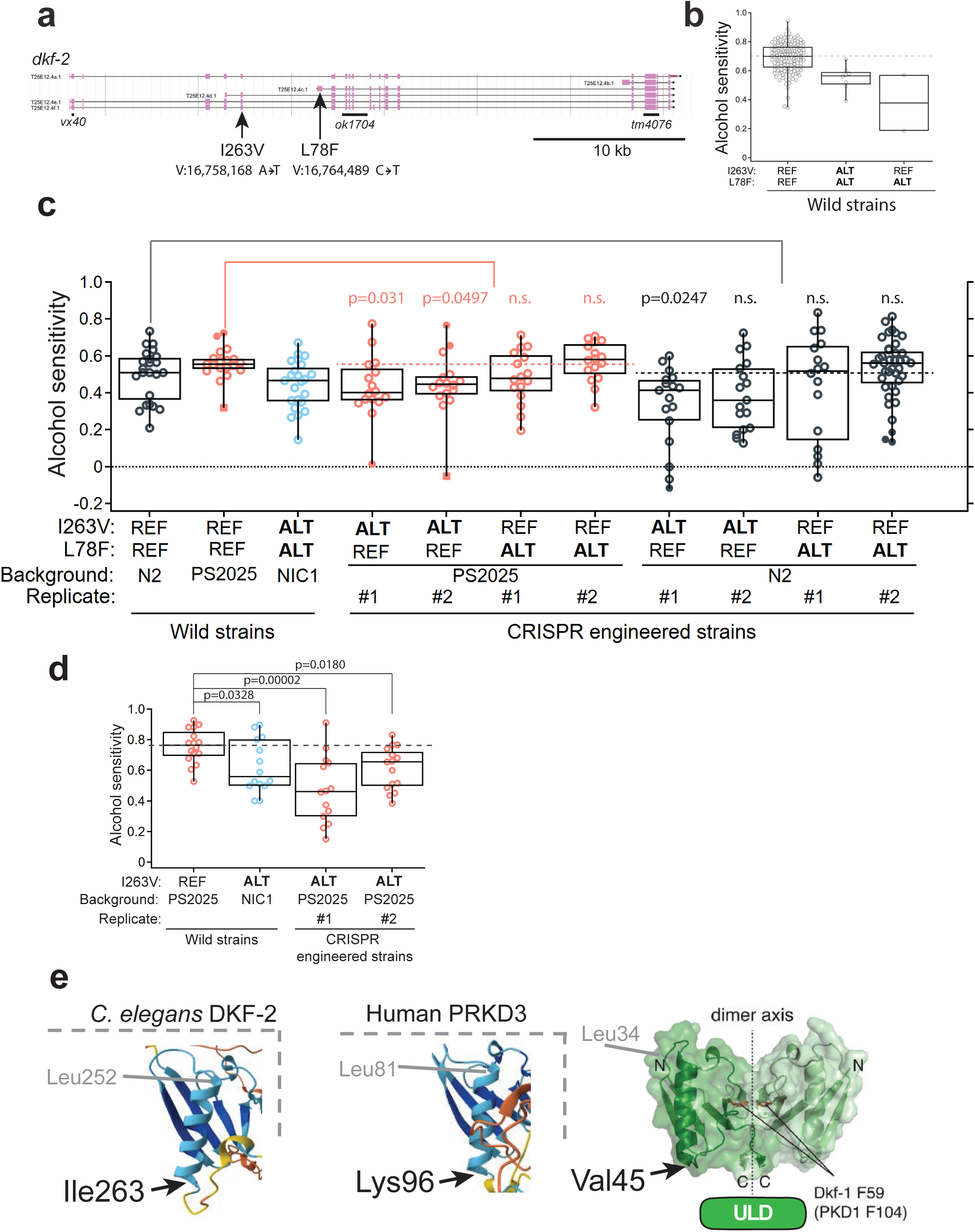
The DKF-2(I263V) variant contributes to alcohol sensitivity. **(a)** Schematic diagram of the *dkf-2* gene locus and location of ALT variants I263V and L78F. **(b)** Alcohol sensitivity plotted for 11 strains that comprise three wild strains and four CRISPR-Cas12a engineered strains made in duplicate. Replacing the ALT variant I263V into PS2025 or N2 backgrounds conferred significant decrease in alcohol sensitivity for three out of four strains, *i.e.,* resistance to intoxication. Replacing the ALT variant L78F had no significant effect in either background. Horizontal colored dashed lines added to help compare to mean of corresponding REF-REF background strains. Planned two-sided t-tests were corrected for multiple comparisons using the Holm–Bonferroni method. Corrected p-values are shown color coded to compare with the corresponding background strain. **(c)** Subset of strains from panel b were retested within a one month period to verify that the ALT variant I263V confers reduced alcohol sensitivity in two replicates of the PS2025 REF background strain (n=15-16). **(d)** Predicted structure of *C. elegans* DKF-2 and human PRKD3 share a single alpha helix and beta sheets in the ubiquitin-like domain (ULD). The I263 residue is in an analogous position at one side of the alpha helix. Crystal structure of *C. elegans* protein kinase D paralog DKF-1 shown for comparison (modified from Reinhardt *et al*., 2020). Box plots show the median, interquartile range, data spread, and potential outliers for each group.

Because the loss-of-function mutants displayed lower alcohol sensitivity relative to strains that harbor the *dkf-2* REF allele, we hypothesized that the candidate causal natural variant(s) in *dkf-2* are hypomorphic, because the ALT allele at the marker likewise covaried with lower alcohol sensitivity. To test whether *dkf-2* plays a role in the alcohol sensitivity of the wild strains and to probe the nature of the candidate causal variants, we generated two independent *dkf-2* deletion alleles in each of two representative wild strain backgrounds: NIC1, which harbors the *dkf-2* ALT alleles, and PS2025, which harbors the *dkf-2* REF alleles. We found that novel deletions in the ALT background did not change alcohol sensitivity, but deletions in the REF background significantly reduced alcohol sensitivity (Fig. 3d).

We next performed a hemizygosity test. We crossed PS2025 and NIC1 wild strains with an N2 background strain carrying the *dkf-2* deletion allele *tm4076*. Interestingly, we found that F_1_ *dkf-2* hemizygotes resembled their respective REF or ALT allele parental phenotypes (Fig. 3e). These results provide evidence that the candidate ALT alleles represent likely causal hypomorphic *dkf-2* natural variants. It is worth noting that, although NIC1 has the same phenotype in the hemizygosity tests (mean = 0.504, Fig. 3e) as in GWAS (mean sensitivity = 0.463), it was more sensitive than previous experiments in the assays with deletion of *dkf-2* (mean sensitivity = 0.623, Fig. 3b). This difference was likely caused by aberrantly high untreated egg-laying measures for this set of assays (mean eggs/hour = 84.2) relative to earlier tests (mean eggs/hour = 37) and relative to PS2025 in the same tests (mean eggs/hour = 67.5). Thus, higher basal untreated-measurements of egg laying before ethanol-treated measurements could have inflated the sensitivity of NIC1 in these assays relative to other experiments reported here. We do not know the reasons for this notable difference, but the results from this experiment demonstrate that deletion of the *dkf-2* gene affects alcohol sensitivity in a background-dependent manner, suggesting that natural ALT alleles in *dkf-2* drive differences in alcohol sensitivity.

To determine whether *dkf-2* alleles exert their effects on alcohol sensitivity by gross differences in ethanol metabolism, we measured internal alcohol concentrations for the lab strain (N2), a *dkf-2* wild strain with REF alleles (PS2025), a *dkf-2* wild strain with ALT alleles (NIC1), as well as two *dkf-2* deletion mutants in each of the REF and ALT backgrounds. We found no differences in internal alcohol concentrations after one hour of ethanol exposure between any of the strains (Supplementary Fig. 6). Moreover, alcohol concentrations were consistent with established measures (Alaimo *et al*., 2012). These results suggest that phenotypic differences in alcohol sensitivity were not caused by differences in ethanol metabolism or uptake. Taken together, we conclude that *dkf-2* is a causal gene that contributes to alcohol sensitivity in *C. elegans*.

### An isoleucine-to-valine variant in DKF-2 contributes to differences in alcohol sensitivity

To investigate specific variants in *dkf-2* that could underlie natural variation in alcohol sensitivity, we considered two that are predicted to have impacts on gene function (Fig. 4a). First, we considered V:16,758,168 that is predicted to cause an ALT missense variant (I263V) in DKF-2 isoforms a, e, and f, as well as a missense variant (I41V) in isoform d. The variant is located three nucleotides from the start of a splice site for these isoforms. Second, we considered V:16,764,489 that is predicted to cause a missense variant L78F in isoform c of DKF-2. Out of the 151 wild strains tested, 10 strains had both the 263V and 78F ALT variants and only two strains had only the 78F ALT variant (Fig. 4b).

We prioritized testing two ALT variants predicted to confer alcohol resistance because resistance is less likely than hypersensitivity to arise from nonspecific effects (Davies and McIntire, 2004). To test if either of the natural ALT variants conferred lower alcohol sensitivity, we generated knock-in alleles for I263V or L78F in N2 and PS2025 background strains. These strains harbor REF variants at both loci and are more sensitive to alcohol compared to our comparison strain NIC1 that has both ALT variants. To control for off-target genome-editing effects, we generated two independent strains for each variant replacement, resulting in eight CRISPR-engineered strains to compare with three wild strain controls (Fig. 4c). We plotted data for each unique strain and statistically compared each CRISPR-engineered strain with its wild background correcting for multiple comparisons (Fig. 4c). We found that for three out of four of the engineered strains the I263V ALT variant conferred significantly decreased alcohol sensitivity compared to their corresponding genetic background (two-sided t-tests: N2 vs. N2 I263V #1, p = 0.0247; N2 vs. N2 I263V #2, p = 0.0850; PS2025 vs. PS2025 I263V #1, p = 0.0306; PS2025 vs. PS2025 I263V #2, p = 0.0497; Fig. 4c). By contrast, we found no significant change in alcohol sensitivity with allele replacement of the L78F variant. To confirm the role for the DKF-2 I263V natural variant in differences in alcohol sensitivity and to attempt to reduce experimental variability due to testing over a prolonged period, we repeated analysis for four strains within a one month period. We found that knock-in of the I263V ALT allele significantly reduced alcohol sensitivity in comparison to the PS2025 background strain (Fig. 4d). Overall, these results provide strong evidence that the DKF-2 I263V natural variant underlies the differences in alcohol sensitivity.

To infer how the I263V ALT variant could alter function of DKF-2 in alcohol sensitivity, we considered conservation of this residue at primary and tertiary protein structure levels. At the primary sequence level, I263 in DKF-2 is poorly conserved with human and mouse protein kinase D2 and D3 (PRKD2 and 3) (Supplementary Fig. 7). However, predicted 3D structures show that I263 is likely located at an analogous position across nematode, mouse, and human orthologs. Specifically, I263 and its human equivalent, K96, both represent the terminal residue in an alpha helix within the ubiquitin-like (ULD) domain (Fig. 4e). This predicted alpha helix structure is supported by the crystal structure (2.2 Å resolution) of the ULD domain from the *C. elegans* PKD paralog DKF-1 (Fig. 4e, Reinhardt *et al*., BioEssays 2020). The ULD helps PKD dimerize, which shifts it to a more active state. Thus, the I263V natural variant could decrease DKF-2 activity by interfering with dimerization.

### Pharmacological probing of *dkf-2* and alcohol sensitivity

Because protein kinases represent druggable targets, we next asked whether DKF-2 affects alcohol sensitivity using a selective protein kinase D inhibitor (CID 755673 – Tocris Bioscience). We chose to assay the laboratory-adapted strain N2 and a deletion mutant *dkf-2(ok1704)* in that genetic background to avoid the potential confounding effects of variation in the metabolisms of wild strains. CID 755673 has been shown to disrupt multiple cellular responses of all three human PKDs *in vitro* (Sharlow *et al*., 2008). Although CID 755673 has not been used in *C. elegans*, previous studies found remarkably conserved functions and activation dynamics for *C. elegans* DKF-2 and human PKDs. For example, DKF-2A exogenously expressed in human HEK293 cells is predictably activated by pharmacological activation of endogenous upstream human PKCs (Feng *et al*., 2007). Likewise, DKF-2A expressed in human cell culture shows robust phosphorylation activity towards human PI4KIIIβ when treated with a PKC activator – a known native target of human PRKD2 (Aicart-Ramos *et al*., 2016). We found that the *dkf-2(ok1704)* deletion mutant showed no significant difference in alcohol response when treated with the inhibitor (two-sided t-test, p = 0.84), whereas the N2 strain displayed a non-significant trend towards a decrease in alcohol sensitivity (Supplementary Fig. 8; two-sided t-test p = 0.07, Cohen’s d: 0.584 suggesting a medium effect). Given the non-significant change in alcohol sensitivity, future *C. elegans* studies might consider optimizing treatment or using a different PKD inhibitor.

## Discussion

By harnessing natural variation in wild strains of *C. elegans*, we identified five QTL associated with variation in alcohol sensitivity. Although we dissected only one of the five QTL identified in the GWAS, statistical fine mapping of the remaining QTL suggests several promising candidate genes underlying these loci that should be investigated in future studies (Fig. 1b). Further, we demonstrated a new role for protein kinase D (*dkf-2*) in alcohol sensitivity. To our knowledge, protein kinase D has eluded identification using classical forward-genetic screens for ethanol sensitivity in the *C. elegans* laboratory-adapted strain N2 – nor has PKD attracted attention in the wider field of alcohol research to date. Only a single lab has considered PKD, not in a role for behavioral sensitivity to alcohol but as a potential therapeutic target in experimental alcoholic pancreatitis (Yuan *et al*., 2021; Yuan *et al*., 2022).

### Protein kinases and alcohol response

Although other protein kinases are known effectors of behavioral responses to alcohol (*e.g*., PKCs (Harris *et al*., 1995; Wallace *et al*., 2007), ERKs (Lee & Messing, 2008), PKAs (Pandey *et al*., 2005), and CAM kinases (Zhang *et al*., 2011)), this study is the first to implicate protein kinase D (PKD). PKDs are serine/threonine protein kinases that affect a wide range of cellular processes (Fu & Rubin, 2011; Ellwanger & Hauser, 2013). Most often, PKDs are downstream effectors of protein kinase C (PKC) and diacylglycerol (DAG) via phospholipase C (PLC) β and γ, which are in turn activated by a variety of neurotransmitters, hormones, and other stressors (Fu & Rubin, 2011; Waldron & Rozengurt, 2003) (Supplementary Fig. 9). Recent studies, however, show that PKDs are sometimes activated via PKC-independent autophosphorylation, but the precise mechanism and functional consequences of this pathway remain obscure (Elsner *et al*., 2019; Reinhardt *et al*., 2020).

Using CRISPR-engineered variant swapping, we found evidence that the I263V natural variant contributed to differential alcohol sensitivity. Inspection of this residue on predicted tertiary structure suggests that it resides in the alpha helix within the ULD. The ULD domain mediates dimerization of PKD upon membrane recruitment (via diacylglycerol), which in turn enables *trans*-autophosphorylation of the activation loop thus enabling kinase activation (Reinhardt *et al*., 2020). This mechanism makes PKD activation concentration- and localization-dependent, offering a regulatory checkpoint. The ULD therefore is a key regulatory domain to switch PKD from an inactive (or low activity) state to an active signaling state. We hypothesize that the I263V ALT variant reduces the function of DKF-2 by interfering with this conserved domain and its associated functions.

As integrators of signal inputs from PKC and DAG, PKDs affect a wide array of downstream cellular processes. These include class II HDAC activity (Parra *et al*., 2005; Dequiedt *et al*., 2005), neuronal function and development (Fu *et al*., 2009; Bisbal *et al*., 2008; Czondor *et al*., 2009), and Golgi and membrane dynamics (Rozengurt, 2011), as well as a conserved role in activation of immune and stress tolerance pathways (Kim *et al*., 2010; Feng *et al*., 2007; Ren *et al*., 2009) (Supplementary Fig.8). PKD could plausibly modify alcohol sensitivity by one or more of these pathways, but immune and stress tolerance pathways represent especially promising candidate mechanisms. In *C. elegans, dkf-2* plays a role in innate immune response, where it promotes PMK-1 (ortholog of p38 MAP kinase) activity, and downregulates *daf-16*, a transcription factor that promotes stress tolerance and lifespan through the Insulin/IGF-1-like signaling (IIS) pathway (Lin *et al*., 1997; Lin *et al*., 2001). Thus, PKD sometimes interacts with the p38 MAPK cascade, a pathway containing *nsy-1* (ortholog of human MAP3K), mutants of which are resistant to the toxic effects of ethanol and other drugs of abuse (*e.g*., methamphetamine, MDMA) (Gomez, 2013; Schreiber & McIntire, 2011).

Given the size of the genomic locus and coding sequence of *dkf-2*, it is puzzling that this gene was not identified in earlier forward genetic screens. In some of the *dkf-2* deletion mutants, particularly the *tm4076* mutant where all isoforms of the gene were predicted to be affected, we found a low penetrance sterility phenotype as well as poor gut health. Spontaneous mutagenesis and natural selection in wild populations, on the other hand, could have produced variants subtle enough to affect essential gene function without causing lethality or sterility (Ossowski *et al*., 2010; Schroeder *et al*., 2017).

### Ecological context of ethanol sensitivity

Wild *C. elegans* inhabit decomposing fruits and vegetation, where microbial fermentation could expose them to variable ethanol concentrations (Samuel *et al*., 2016). Reduced sensitivity might help worms maintain feeding, movement, and egg laying in ethanol-rich patches. Greater sensitivity could instead favor avoidance of concentrations that impair these behaviors. Ethanol may also be useful at low concentrations, as it prolongs survival of first-larval and dauer stage *C. elegans* in the absence of bacterial food in laboratory experiments (Castro *et al*., 2012; Artyukhin *et al*., 2015; Kaptan *et al*., 2020). These possibilities suggest that the fitness effects of an ethanol-response variant may depend on the substrate, its microbes, and the worm’s life stage.

Our previous study using the laboratory N2 background found a trade-off between acute ethanol sensitivity and withdrawal severity (Scott *et al*., 2017). *slo-1* loss-of-function mutants resisted acute intoxication but showed more severe withdrawal, whereas gain-of-function mutants were more sensitive to intoxication but showed milder withdrawal. Thus, any advantage of reduced acute sensitivity during prolonged ethanol exposure might be offset by more severe withdrawal. Future studies should investigate whether this potential tradeoff is found for wild strains characterized in this study.

Davies *et al*. (2004) initially interpreted differences between *npr-1* alleles as evidence that natural genetic variation shapes ethanol responses. Subsequent work showed that the NPR-1(215V) allele in the reference strain N2 arose during laboratory domestication and was favored under laboratory cultivation, whereas wild strains generally carry the ancestral 215F allele (Sterken *et al*., 2015). Thus, although *npr-1* affects ethanol responses, the original allele comparison does not establish that ethanol sensitivity was selected in natural habitats. Whether the variation identified in the 151 wild strains here reflects local adaptation to ethanol or associated effects on other traits remains to be tested under ecologically realistic conditions.

### Conclusions and limitations

Here, we used a high-throughput behavioral screen to measure acute behavioral sensitivity to ethanol intoxication in a diverse set of over 150 wild strains of the nematode *Caenorhabditis elegans*. Using GWAS, we identified five loci associated with alcohol sensitivity. Next, we causally validated the role of a highly conserved orthologue of PKD in *C. elegans* alcohol response. We further find that a PKD inhibitor trends towards affecting alcohol intoxication sensitivity in a *dkf-2*-dependent manner. Together, these data reveal a new role for PKD in alcohol intoxication, a trait that significantly predicts later life alcohol abuse in human populations. If future studies demonstrate a conserved role for PKD in mammalian alcohol response, these findings represent an exciting potential avenue for novel AUD treatments.

Already, work in mammals has shown that PKCs are druggable targets with substantial effects on AUD traits (Lee & Messing, 2008). However, relative to PKCs, PKDs were only recently discovered (Fu & Rubin, 2011), thus, far fewer studies and available reagents are available for researchers to exploit. Future work should aim to tease apart the mechanisms by which PKD affects sensitivity to ethanol intoxication.

Lastly, we have provided evidence using genome replacements and analyses of homozygous and hemizygous deletions to support a role for *dkf-2* natural variation in modulating alcohol sensitivity. A complete hemizygosity analysis will be needed to exclude background effects. We acknowledge that the *dkf-2* could harbor variants in addition to I263V that work together to explain the observed natural differences in alcohol sensitivity (*e.g*., L78F in certain circumstances). Identifying causal variants might reveal conserved mechanisms relevant to alcohol use disorder risk and illuminate how *C. elegans* has adapted to ethanol in its environment.

## Data availability

The authors affirm that all data necessary for confirming the conclusions of the article are present within the article, figures, and supplementary data file. Strains and plasmids are available upon request.

## Acknowledgments

For strains and public data, we thank the *Caenorhabditis* Genetics Center, which is funded by NIH (P40 OD010440), WORMBASE, which is funded by NIH (U41 HG002223), and CaeNDR, which is funded by NIH (R50ES037948). We appreciate the Pierce and Andersen labs for advice, Nicole Liachko for the *dkf-2(ok1704)* strain, and Susan Rozmiarek and Cory Gentry for technical assistance.

## Author contributions

B.C., E.C.A., and J.T.P. designed research; B.C., T.C.O., B.V.K., and J.T.P. performed research; M.K.M. performed mapping; S.J.K. contributed reagents; B.C., E.C.A., and J.T.P. analyzed data; J.T.P. and B.C. acquired funding; and B.C. wrote first draft, B.C., E.C.A., and J.T.P. worked on final paper.

