## Supplemental Info for "Natural variation in protein kinase D modifies alcohol sensitivity in *Caenorhabditis elegans*"

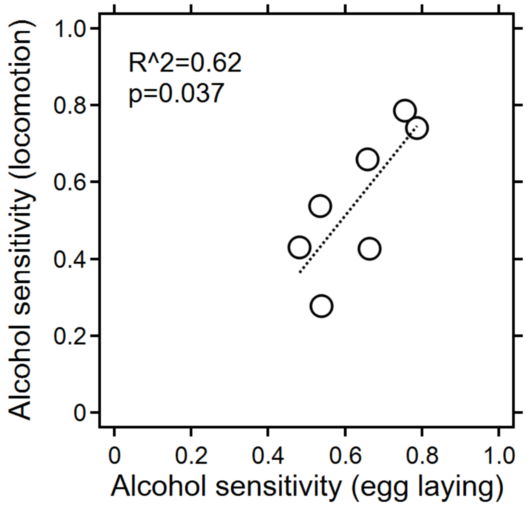


**Supplementary Figure 1. Correlation between alcohol induced depression in egg-laying and locomotion.** Wild *C. elegans* strains were assayed for both alcohol sensivitity as determined by alcohol-induced depression in locomotion and egg-laying. The two measures are significantly correlated ($R^2$=0.62, p=0.037). Raw data found in Supplementary data.

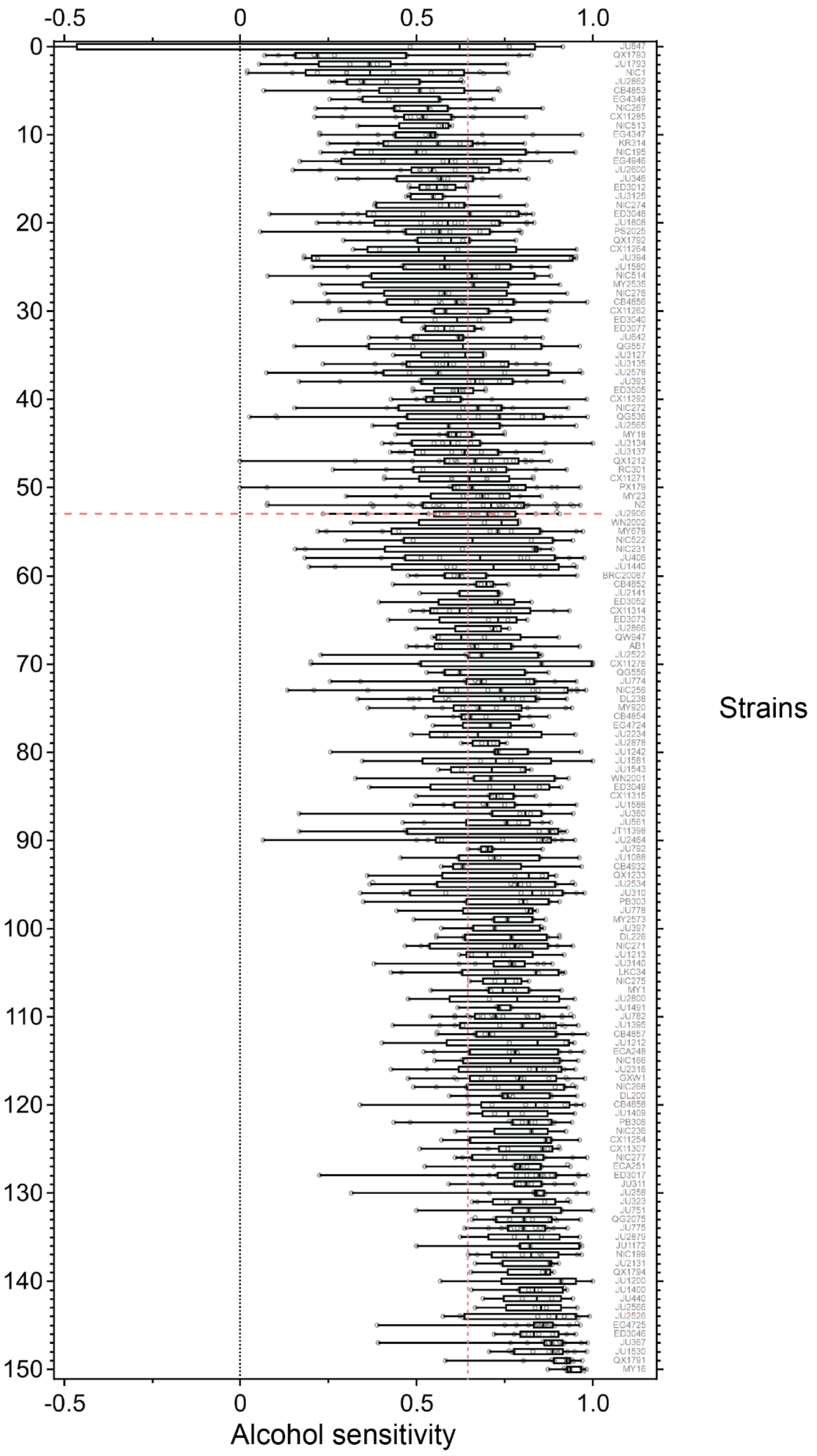
-0.5
0
0.5
1.0
0
10
20
30
40
50
60
70
80
90
100
110
120
130
140
150
Alcohol sensitivity
Strains

**(Previous page) Supplementary Figure 2. Correlation between alcohol induced depression in egg-laying and locomotion.** Wild *C. elegans* strains (N=151) were assayed for alcohol sensivitity as determined by egg-laying. Box plots show the median, interquartile range, data spread, and potential outliers for each group. N2 data noted by red dashed lines. Raw data found in Supplementary data.

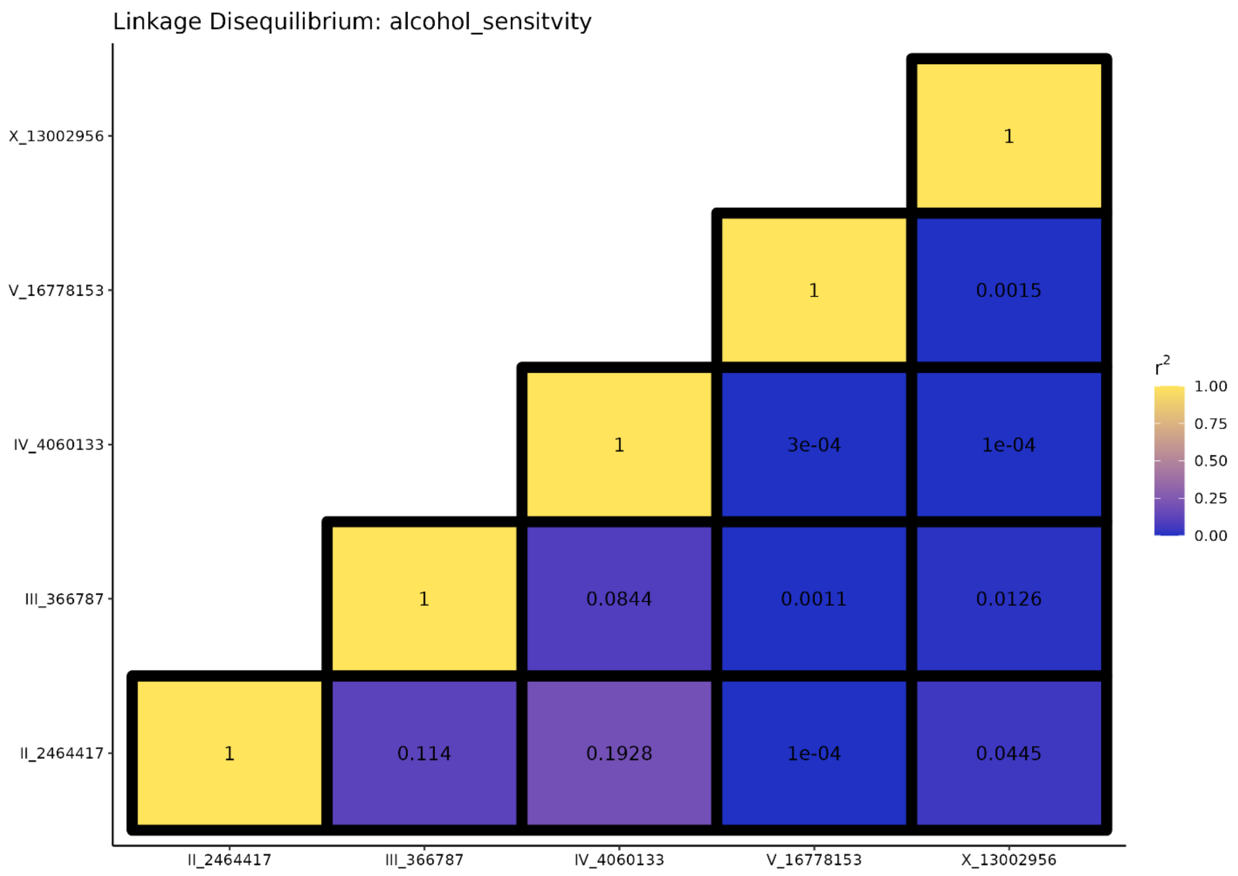


**Supplementary Figure 3. Linkage disequilibrium between significant QTL.** Linkage disequilibrium ($r^2$) between each of the five significant QTL is shown

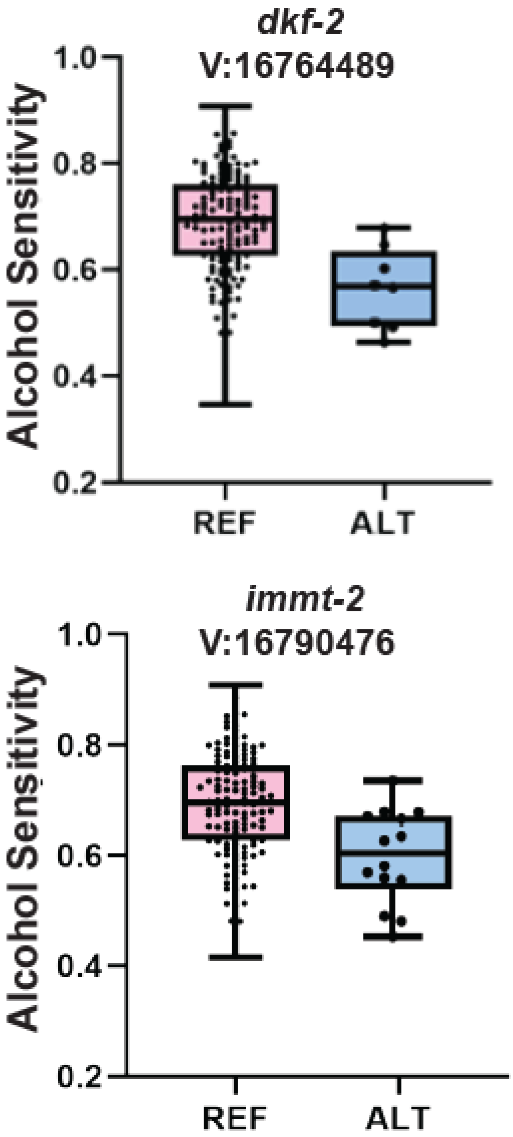


**Supplementary Figure 4. Fine mapping chromosome V QTL associated with decreased sensitivity.** Each point in boxplots represents the average alcohol sensitivity of a single strain. Y-axis denotes alcohol intoxication sensitivity, and genotype at SNV is shown along the X-axis (REF or ALT). Average phenotypes of all strains were segregated by their genotype at the denoted allele. Top and bottom graphs show phenotypes segregated by genotype at two impact mutation candidate variants identified by fine mapping.

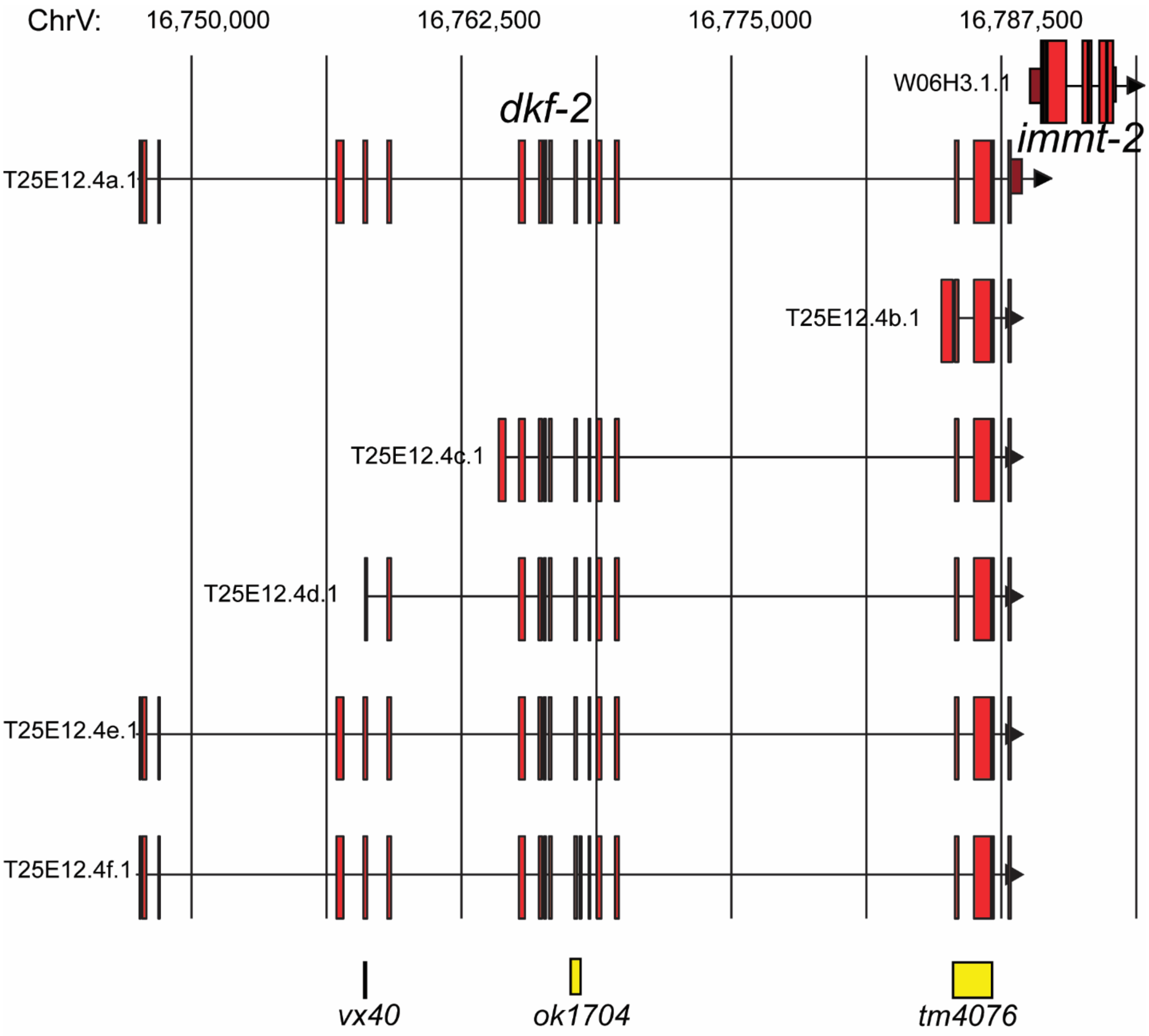


**Supplementary Figure 5. Approximate position of N2 background deletion mutants in *dkf-2* splice isoforms.** Genomic loci of each of the six *dkf-2* isoforms are diagrammed above. Red boxes indicate exons, and solid lines connecting boxes indicate introns. Each of the three N2 background deletion mutants are shown using yellow boxes below.

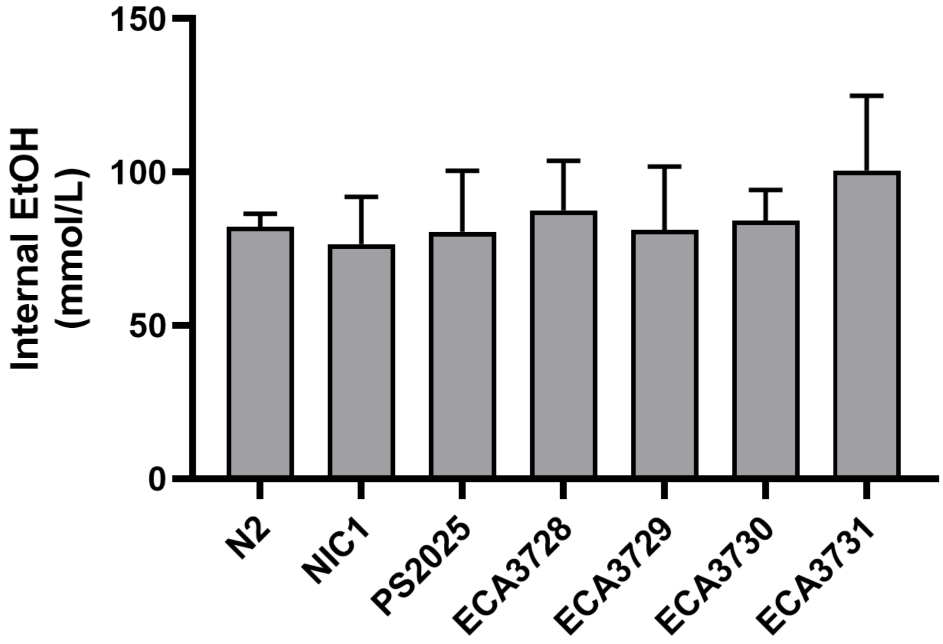


**Supplementary Figure 6. Internal ethanol concentrations did not differ with respect to *dkf-2* alleles.** Internal ethanol concentrations were measured for the N2 lab strain (mean = 82.24 mmol/L), NIC1 ALT background wild strain (mean=76.52 mmol/L; SEM=15.42; n=4), PS2025 REF background wild strain (mean = 80.59 mmol/L; SEM=19.8; n=3), and for two independent CRISPR-generated deletions in each of the wild-strain backgrounds – for PS2025 background ECA3729 (mean=81.14 mmol/L; SEM=20.7; n=4) and ECA3730 (mean=84.19 mmol/L; SEM=10.03; n=4) - for NIC1 background ECA3728 (mean= 87.53 mmol/L; SEM=16.2; n=4) and ECA3731 (mean=100.61 mmol/L; SEM=24.3; n=3). No significant differences were found between strains (ANOVA; F=0.2024; DF=25; p=0.9719).

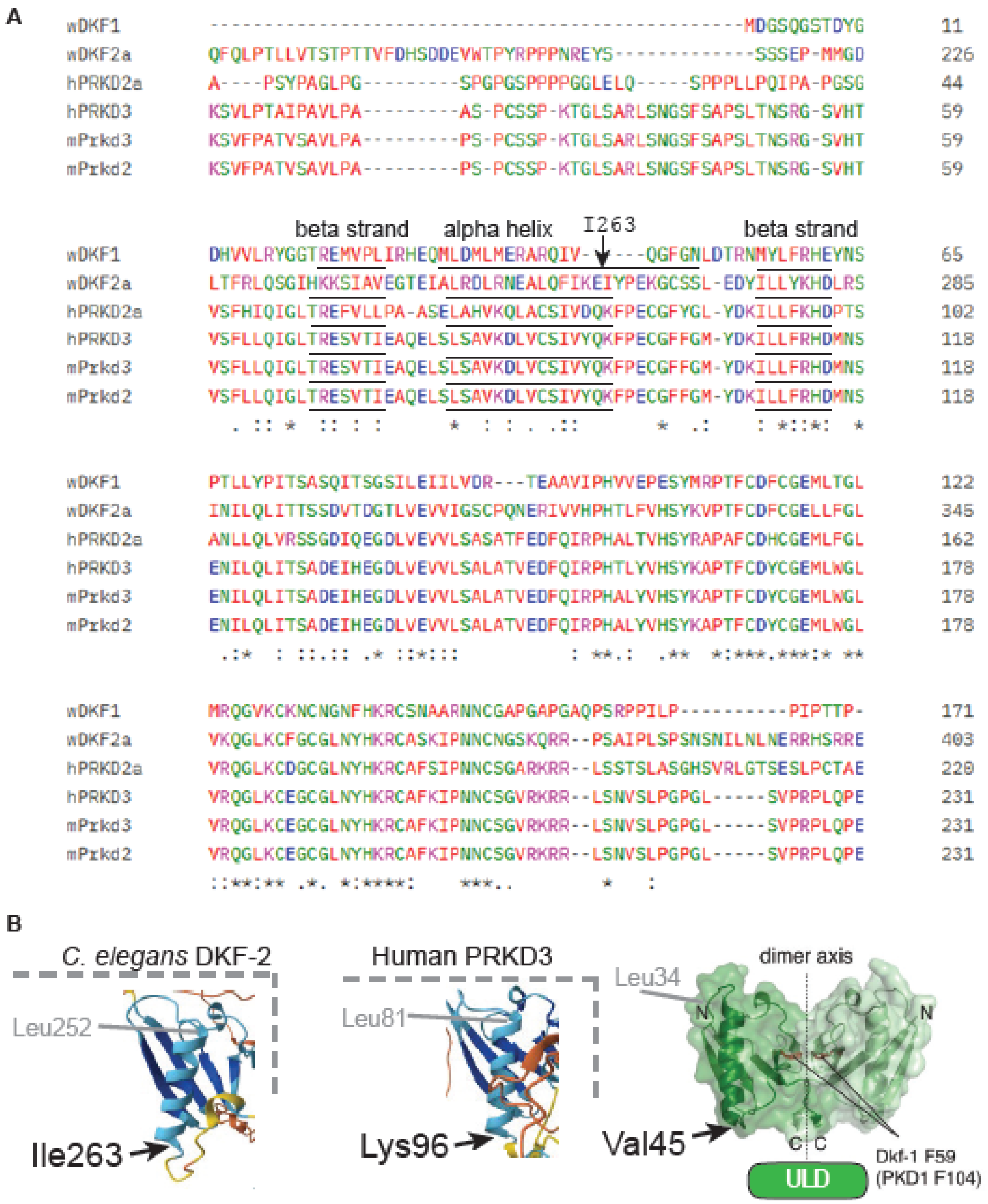


**Supplementary Figure 7. Alignment of conserved protein kinase D proteins. A)** Primary sequence of amino acids aligned by Clustal Omega and color-coded using Clustal X scheme, which colors residues based on their physiochemical properties (Sievers and Higgins, 2011). Alpha helix region and adjacent strands of beta sheets of ULD are underlined as determined by Alphafold2 or crystal structure (Jumper et al., 2022; Reinhardt et al. 2020). I263 residue highlighted with arrow. **B)** ULD structure predicted by Alphafold2 (DKF-2 and PRKD3) or crystal structure (DKF-1) modified from Reinhardt et al., (2020). 3D structures in panel B reproduced from Figure 4C for comparison with primary sequence in panel A.

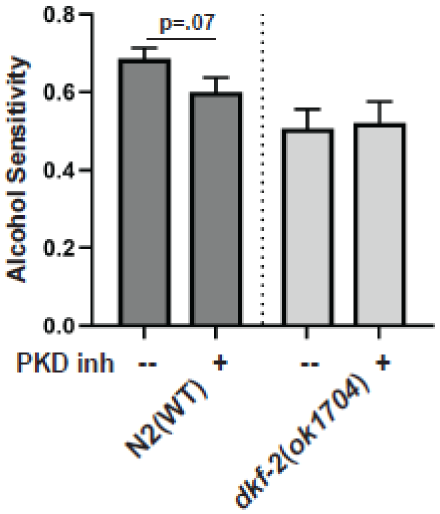


**Supplementary Figure 8. Testing alcohol sensitivity with protein kinase D inhibitor.** Treatment with a PKD inhibitor (CID 755673) trends towards lower alcohol sensitivity in the N2 lab strain (mean & SEM, n= 18-20, two-sided t-test p=0.07, Cohen's d: 0.584 -medium effect) but shows no effect on alcohol sensitivity in a *dkf-2* deletion mutant allele *ok1704*.

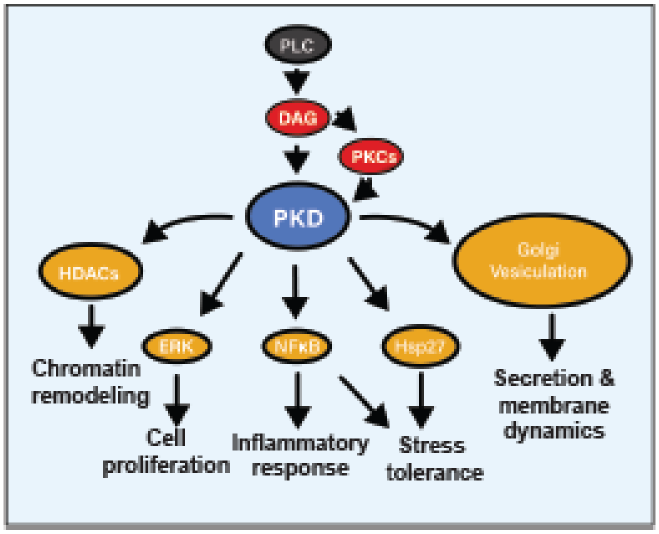


**Supplementary Figure 9. Known interactions of PKD.** PKDs are most often downstream effectors of DAG-PKC signaling cascades. Activated PKDs mediate a wide array of cellular processes *e.g.*, chromatin dynamics, cell growth and proliferation, innate immunity and inflammatory response, general stress tolerance and Golgi vesiculation. Recent studies show that PKD can activate via PKC-independent autophosphorylation of the conserved ULD domain.

.

**Supplementary Table 1. GWA summary statistics for each of the five QTL**

| CHROM | marker | log10p | startPOS | peakPOS | endPOS |
|---|---|---|---|---|---|
| II | II:2464417 | 5.691478 | 1284204 | 2464417 | 2776923 |
| III | III:366787 | 5.163477 | 12249 | 366787 | 1618298 |
| IV | IV:4060133 | 5.397415 | 2671608 | 4060133 | 12898625 |
| V | V:16778153 | 4.425160 | 16616122 | 16778153 | 17602325 |
| X | X:13002956 | 4.404005 | 11187104 | 13002956 | 14806635 |
